# Chlorophyllide *a* oxygenase (CAO) gene duplication across the Viridiplantae

**DOI:** 10.1101/2025.02.10.637544

**Authors:** Mackenzie C. Poirier, Roberta Wright, Marina Cvetkovska

## Abstract

Viridiplantae, a diverse group of green plants and alga that have evolved from a common ancestor, are unified in their ability to produce and use two types of chlorophyll (chlorophyll *a* and chlorophyll *b*) to capture light energy. In addition to playing a role in light harvesting, chlorophyll *b* is required at the appropriate level for the accumulation, assembly, and stability of light harvesting complexes within the photosynthetic apparatus. Chlorophyll *b* is synthesized from chlorophyll *a* by the enzyme chlorophyllide *a* oxygenase (CAO), a Rieske-type mononuclear non-heme iron oxygenase. A regulatory degron sequence, described in detail only in land plants, regulates the stability of CAO proteins based on the availability of chlorophyll *b.* Recent identification of CAO gene duplication in bryophyte and green algal species, combined with expanded availability of sequenced genomes within the Viridiplantae, prompted further investigation into the role of gene duplication in the evolution of chlorophyll *b* biosynthesis. Examination of genomes from 246 plant and algae species revealed independently occurring CAO duplications throughout the Viridiplantae, with a higher prevalence of duplication in land plants compared to their algal relatives. Additionally, we demonstrate that the degron sequence is poorly conserved in chlorophytes, but first appears as a conserved sequence in charophytes, and is very highly conserved among the embryophytes. The evolutionary history and functional role of CAO throughout the Viridiplantae lineage is discussed based on these key observations, adding to our understanding of chlorophyll *b* biosynthesis and the role of CAO in photosynthetic species.

## Introduction

Photosynthetic organisms form the base of most food chains and provide the organic carbon compounds that support life in our biosphere. The capture of light energy by chlorophyll is the first step in the process of photosynthesis that fixes atmospheric CO2 into stable organic compounds. Overwhelming evidence indicates that oxygenic eukaryotic photosynthesis that occurs in the chloroplasts of modern-day plants and algae traces its origins from an endosymbiotic event with cyanobacteria-like prokaryotes (Blankenship 2010; Cardona 2019; Sánchez-Baracaldo and Cardona 2020). Viridiplantae (or green plants) have primary chloroplasts derived from an ancient endosymbiosis, and encompass two major clades: the chlorophytes and the streptophytes.

Chlorophytes are a monophyletic group of marine, freshwater, and terrestrial green algae. With ∼8,000 described species (Guiry 2024), this group encompasses a large diversity of adaptations, morphologies, and life histories (Leliaert et al. 2012). Streptophyta includes the charophytes, ∼5,500 species of largely freshwater green algae (Guiry 2024) and the embryophytes, or land plants. Land plants are thought to have diverged from a single clade within the streptophytes (de Vries and Archibald 2018) and number ∼430,000 species (Ruggiero et al. 2015). One characteristic that unifies members of the Viridiplantae is their ability to synthesize and use two types of chlorophylls: chlorophyll *a* (Chl *a*) and chlorophyll *b* (Chl *b*).

The photosynthetic apparatus in the Viridiplatae is derived from the ancestral cyanobacterial endosymbiont, which used chlorophyll as the main pigment in their photosystems (Xiong and Bauer 2002; Sánchez-Baracaldo and Cardona 2020). Chl *a* is ubiquitously present in all oxygenic photosynthetic eukaryotes, and functions in both energy capture in the antenna light harvesting complexes (LHCs) and in driving electron transfer in the photosystem II (PSII) and photosystem I (PSI) reaction centers. Chl *b* is specific to the Viridiplantae although it has also been detected in prochlorophytes and *Acaryochloris*, unique photosynthetic prokaryotic groups that lack typical cyanobacterial phycobilin light-harvesting pigments (Palenik and Haselkorn 1992; Roche et al. 1996; Tomitani et al. 1999; Partensky et al. 2018). Chl *b* resides predominantly in the peripheral LHCs (Neilson and Durnford 2010) although it has been detected in the core complexes of some deep water marine chlorophytes (Kunugi et al. 2016).

Strict Chl *a/b* stoichiometry is required for optimal energy transfer during photosynthesis. Maintaining the correct Chl *a/b* ratios is a dynamic process and indicative of adaptation to different light conditions (Tanaka and Tanaka 2007). In addition to playing a role in light harvesting, Chl *b* is required at appropriate levels for the accumulation, assembly, and stability of LHCs in algae (Bujaldon et al. 2017), bryophytes (Zhang et al. 2023), and angiosperms (Król et al. 1995; Reinbothe et al. 2006; Kim et al. 2009; Nick et al. 2013). Under low light conditions, the amount of Chl *b* relative to Chl *a* increases, leading to larger LHC antenna size, which maximizes the surface area for light absorption (Kunugi et al. 2016; Kume et al. 2018; Ueno et al. 2019). Thus, the amount of Chl *b* is a major regulator of light-harvesting capacity.

Chlorophyll biosynthesis and turnover occurs through a complex multistep pathway (reviewed in Brzezowski et al. 2015; Willows 2020). Chl *b* is synthesized from Chl *a*, via the intermediate 7- hydromethyl chlorophyll *a*, by the action of a single enzyme: chlorophyllide *a* oxygenase (CAO) (Tanaka et al. 1998; Espineda et al. 1999; Oster et al. 2000; Mueller et al. 2012). Chl *b* can be converted back into Chl *a* by chlorophyll *b* reductase (CBR) and 7-hydroxymethyl reductase (HCAR) to complete the cycle (Figure 1; reviewed in detail in Tanaka and Tanaka, 2019). The chlorophyll cycle plays an essential role in many biological processes, including biogenesis of LHCs, antenna size regulation during light acclimation, and chlorophyll degradation during senescence. While many of the intricacies of how this pathway is regulated are still not fully understood (Tanaka and Tanaka 2019), it has been suggested that CAO levels and activity are key determinants of Chl *b* accumulation.

**Figure 1.**
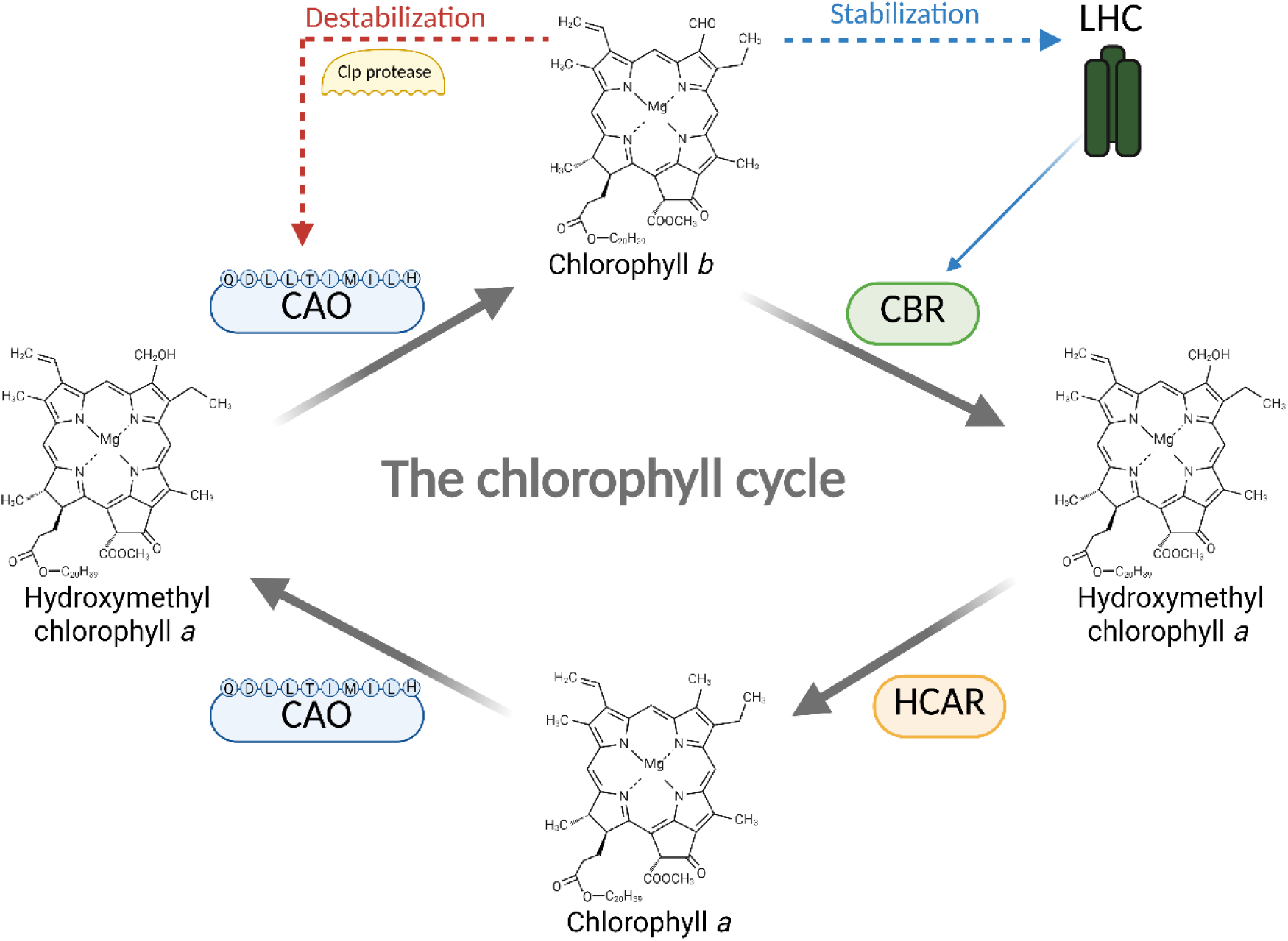
The chlorophyll cycle in land plants. Chlorophyll *a* is converted to chlorophyll *b* via the intermediate 7-hydromethyl chlorophyll *a*, by the action of the enzyme chlorophyllide *a* oxygenase (CAO). In a negative feedback loop, high levels of chlorophyll *b* trigger the binding of a stromal Clp protease to the degron motif (QDLLTIMILH) and the subsequent destabilization of CAO (dashed red line). Chlorophyll *b* is necessary for the stabilization of Light Harvesting Complexes (LHC) in plants (dashed blue line). Excessive amounts of LHC or accumulation of energetically uncoupled non-functional LHCs (e.g., during senescence), triggers the accumulation of Chlorophyll *b* reductase (CBR; blue line) and the conversion of chlorophyll *b* to chlorophyll *a* via the action of CBR and 7- hydroxymethyl reductase (HCAR). This regulatory mechanism of CAO and CBR enables the fine-tuning and optimization of chlorophyll *b* and LHC levels (adapted from Tanaka and Tanaka 2019).

CAO is part of the Rieske-type mononuclear non-heme iron oxygenase group of proteins (Gray et al. 2004). It catalyzes the conversion of the methyl group of Chl *a* into a formyl group to form Chl *b* through two oxygenation reactions, likely using ferredoxin as a reductant (Oster et al. 2000). CAO has three main regions: an N-terminal regulatory domain and a C-terminal catalytic domain, which are connected by a more variable linker region (Nagata et al. 2004). The catalytic domain contains a Rieske cluster and a mononuclear iron-binding domain, both of which are necessary for the enzymatic activity of CAO (Yamasato et al. 2005; Kunugi et al. 2013). The regulatory domain has been characterized in detail only in land plants. In *Arabidopsis*, a conserved sequence (QDLLTIMILH) termed a degron was demonstrated to regulate Chl *b* accumulation by affecting CAO protein turnover. In a negative feedback loop, high levels of Chl *b* trigger the binding of a stromal Clp protease to the degron motif leading to destabilization of the CAO protein; this, in turn, halts Chl *b* biosynthesis (Sakuraba et al., 2009; Nakagawara et al., 2007).

The evolutionary origins of CAO can be traced back to prochlorophytes, which encode a CAO gene homolog (Tomitani et al. 1999; Nagata et al. 2004). Notably, the prokaryotic CAO-like enzymes lack the regulatory N-terminal domain, suggesting that the Chl *b*-dependent mechanism for controlling CAO stability was acquired in eukaryotes. Recent work on 9 chlorophyte, 2 charophyte, and 3 embryophyte species suggested that the degron recognition sequence first appeared in charophytes as an adaptation to high-light environments in shallow water (Kunugi et al. 2016).

These conclusions, however, were based on a very limited sample of representative species and the presence of a degron in the CAO sequence in charophytes was inferred based on a single species (*Klebsormidinium flaccidum*). This hypothesis requires further validation.

The CAO protein is encoded by a single gene in most plants and algae (Kunugi et al. 2016; Schumacher et al. 2022), with only a very few reported exceptions. Two CAO genes were detected in rice (Lee et al. 2005) although only one of them is necessary for Chl *b* synthesis (Jung et al. 2021). Recent work revealed CAO duplication in the bryophyte *Physcomitrium patens,* where both genes contribute to Chl *b* accumulation (Zhang et al. 2023). CAO duplication was also reported in two Antarctic *Chlamydomonas* species, where the CAO paralogs in each species appear to be derived from independent duplications events rather than an ancestral one (Cvetkovska et al. 2019; Poirier et al. 2025).

These insights prompted us to further examine the *CAO* gene content and the presence of the regulatory domain across the diverse plant and algal groups. A systematic examination for the presence of *CAO* across the Viridiplantae has not yet been preformed. In this work we took advantage of the recent increase in publicly available plant and algal genomes to examine the occurrence of CAO genes in diverse species across Viridiplantae and demonstrate a widespread occurrence of CAO duplicates in this group. Furthermore, building on the work by Kunugi et al. (2016), we have examined the occurrence of a degron sequence in 326 unique CAO sequences in algae and plants. We demonstrate that this regulatory region is poorly conserved in chlorophytes, compared to charophyte and embryophyte CAO sequences, suggesting a different mechanism for Chl *b* regulation in early aquatic lineages.

## Materials and Methods

### CAO sequence identification and analyses

To identify *CAO* sequences and duplication events with high accuracy, we used publicly available, high quality, and fully annotated genomes. Chlorophyte and streptophyte genomes were retrieved from Phycocosm (Grigoriev et al. 2021), while embryophyte genomes were obtained from Phytozome v13 (Goodstein et al. 2012). Putative *CAO* genes were identified with a tBLASTn search with the *Chlamydomonas reinhardtii* (CrCAO; Cre01.g043350) and the *Arabidopsis thaliana* (AtCAO; AT1G44446) full-length CAO peptides (Merchant et al. 2007; Lamesch et al. 2012). A strict e-value cutoff of ≤e-30 was taken as a threshold, to avoid many unspecific hits due to the conserved nature of the Fe-binding and Rieske cluster domains present among diverse plant and algal proteins (Schmidt and Shaw 2001; Ferraro et al. 2005; Przybyla-Toscano et al. 2021). Putative *CAO* sequences were manually curated and truncated or low-quality sequences were excluded from downstream analyses. Curated *CAO* sequences were aligned using MUSCLE (Edgar 2004).

The degron sequence (QDLLTUMILH) was identified by multiple sequence alignment of the full-length CAO amino acid sequences and the regulatory N-terminal domain. In accordance with Sakuraba et al. 2009, we identify the degron as “highly conserved” if 8 out of the 10 amino acids are perfectly conserved, “moderately conserved” if 4 to 8 amino acids are conserved, and “poorly conserved” if less than 4 amino acids are conserved. Sequence logos for each major group (chlorophyte algae, charophyte algae, non-vascular plants and vascular plants) were generated by Geneious Prime 2024.11 (Dotmatics), where the size of the letter reflects its frequency. To examine the expression of CAO duplicate genes, we screened previously published transcriptomes from 16 species with representatives from the Chlorophytes (Arriola et al. 2018), Charophytes (Cheng et al. 2019), non-vascular (Perroud et al. 2018; Healey et al. 2023) and vascular plants (Zhu et al. 2019; Wang et al. 2021; Liu et al. 2022b; Shang et al. 2023; Guo et al. 2025; He et al. 2025; Mascuñano et al. 2025; Roy et al. 2025). Only transcriptomes that report FPKM, RPKM and TPM values obtained from organisms cultivated at optimal conditions were considered (Supplementary Table S4).

### Phylogenetic Inference

The analyses described above resulted in 374 individual CAO sequences (Supplementary Table S1). To avoid inaccurate inference and over-representation, we manually removed heterodimeric CAO proteins, where the Rieske domain and the Fe-binding domain are located on different genes (Kunugi et al. 2013) (Supplementary Table S1). We also removed identical CAO sequences isolated from algal cultures identified as the same species but belonging to different strains or culture collections (*Scenedesmus obliquus*, *Auxenochlorella protothecoides*, *Chlorella sorokiniana*, *Ostreococcus tauri*, *Mesostigma viride*, *Zygnema circumcarinatum*). This resulted in a 280 amino acid alignment of 326 unique CAO sequences across 246 species.

These manually curated CAO genes were translated into amino acid sequences, aligned with MUSCLE and trimmed to remove gaps and ambiguously aligned regions with Gblock (Castresana 2000; Talavera and Castresana 2007). Maximum likelihood trees were inferred in the CIPRES Science Gateway (Miller et al. 2015) using RAxML v8.0 (Stamatakis 2014) with 1000 bootstraps with the Whelan Goldman matrix for globular proteins (WAG), a gamma shape parameter, and empirical estimation of invariable sites. The trees were rooted using prokaryotic CAO from *Acaryochloris thomasi* (WP_110987895.1) and *Prochlorotrix hollandica* (WP_017713323.1) as an outgroup.

Tandem duplications were defined as those located on the same chromosome or contig within <10 genes of each other, based on the genomic coordinates retrieved from Phytozome v13 or Phycocosm. To identify ancestral and lineage-specific duplications, we used Notung v2.9 (Stolzer et al. 2012) and reconciled the CAO trees with a reference species tree derived from NCBI taxonomy (Schoch et al. 2020) via the PhyloT v2 tool (https://phylot.biobyte.de). Putative ancestral duplications were only reported when occurring on branches with bootstrap support ≥85%. All trees were visualized in iTOL v6.8.2 (Letunic and Bork 2021). Previously inferred taxonomic information for chlorophytes and streptophytes was confirmed via AlgaeBase, University of Galway (www.algaebase.org), and for land plants via the Plants Of the World Online, Royal Botanic Gardens, Kew (https://powo.science.kew.org).

## Results and Discussion

### Occurrences of multiple CAO genes are widespread among the Viridiplantae

Our analysis demonstrates a widespread occurrence of *CAO* gene duplication across the Viridiplantae. Multiple gene copies are more prevalent among the angiosperms, where ∼41% of examined species encode for more than one *CAO* gene. In comparison, we detected two *CAO* gene copies in only ∼11% of chlorophyte and ∼20% of charophyte species (Table 1; Table 2, Supplementary Table S1; Supplementary Table S3). All 23 species where we detect more than two *CAO* genes are members of the Embryophyta. The modern-day cultivar of sugarcane (*Saccharum* sp. R570) encodes for 6 *CAO* copies, the largest number we report. This species is a result of inter- specific hybridization (*Saccharum officinarum* x *spontaneum;* Dumont et al. 2022), which has resulted in a very large and highly redundant genome (∼0.3 Gb, ∼48,000 genes) (Healey et al. 2024). It must be noted that these results are likely affected by the availability of many more angiosperm genomes in public databases, while other plant and algal groups are not as well represented. As more genomes are sequenced, these analyses will have to be revisited.

**Table 1:**
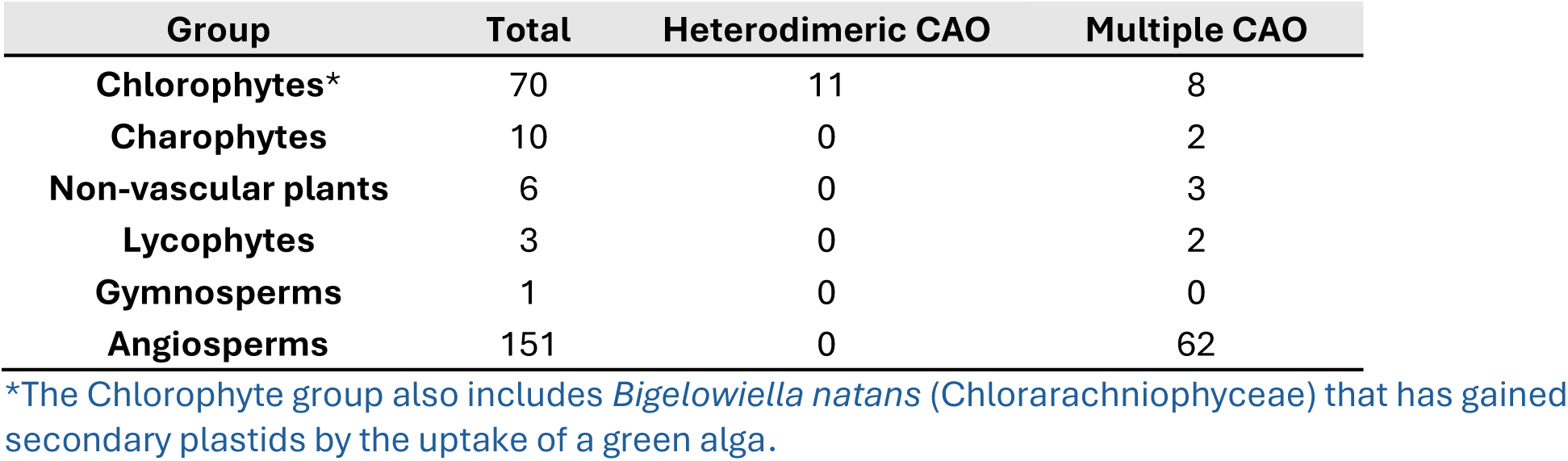
Number of species across the Viridiplantae with fully sequenced and annotated genomes available through Phytozome (Total), compared to the number of species that encode for multiple *CAO* genes (Multiple CAO) and number of species that encode for two *CAO* genes that form heterodimeric complex (Heterodimeric CAO).

| Group | Total | Heterodimeric CAO | Multiple CAO |
| --- | --- | --- | --- |
| **Chlorophytes\*** | 70 | 11 | 8 |
| **Charophytes** | 10 | 0 | 2 |
| **Non-vascular plants** | 6 | 0 | 3 |
| **Lycophytes** | 3 | 0 | 2 |
| **Gymnosperms** | 1 | 0 | 0 |
| **Angiosperms** | 151 | 0 | 62 |

\*The Chlorophyte group also includes *Bigeloviella natans* (Chlorarachniophyceae) that has gained secondary plastids by the uptake of a green alga.

**Table 2:** Distribution of *CAO* duplicate genes across Viridiplantae. Only genes with a confirmed Riske conserved cluster, Fe-binding domain (all species), and degron sequence (Streptophytes only) are listed. A checkmark (✓) indicates that the CAO genes are positioned in a tandem on the same chromosome within the genome, while a cross (x) indicates the genes are located on different chromosomes. Accession numbers can be found in Supplementary Table 1 (continued on next page).

| Taxon | Species | #CAO genes | Tandem |
| --- | --- | --- | --- |
| **Chlorophyta** |  |  |  |
| Chlorophyceae | *Chlamydomonas priscui* | 2 | x |
| Chlorophyceae | *Volvox reticuliferus* | 2 | ✓ |
| Chlorophyceae | *Tetradismus deserticola* | 2 | x |
| Trebouxiophyceae | *Chlorella* sp. A99 | 2 | x |
| Trebouxiophyceae | *Chlorella sorokiniana* | 2 | x |
| Trebouxiophyceae | *Chlorellaceae* sp. | 2 | x |
| Trebouxiophyceae | *Micractinium conductrix* | 2 | x |
| Pedinophyceae | *Pedinomonas minor* | 2 | x |
| **Charophyta** |  |  |  |
| Zygnematophyceae | *Mesotaenium endlicherianum* | 2 | x |
| Zygnematophyceae | *Spirogloea muscicola* | 2 | x |
| **Bryophyta** |  |  |  |
| Bryopsida | *Physcomitrium patens* | 2 | x |
| Sphagnopsida | *Sphagnum fallax* | 2 | x |
| Sphagnopsida | *Sphagnum magellanicum* | 3 | x |
| **Lycophyta** |  |  |  |
| Equisetopsida | *Diphasiastrum complanatum* | 4 | ✓ |
| Equisetopsida | *Ceratopteris richardii* | 2 | x |
| **Angiosperms** |  |  |  |
| Laurales | *Cinnamomum kanehirae* | 2 | ✓ |
| Nymphaeales | *Nymphaea colorata* | 2 | x |
| Asparagales | *Agave tequilana* | 2 | x |
| Asparagales | *Yucca aloifolia* | 3 | x |
| Asparagales | *Yucca filamentosa* | 3 | x |
| Zingiberales | *Musa acuminata* v1 | 2 | x |
| Alismatales | *Spirodela polyrhiza* v2 | 2 | x |
| Poales | *Typha latifolia* | 2 | x |
| Poales | *Eleusine coracana* | 2 | x |
| Poales | *Oryza sativa* | 2 | ✓ |
| Poales | *Thinopyrum intermedium* | 3 | x |
| Poales | *Brachypodium hybridum* | 2 | x |
| Poales | *Brachypodium mexicanum* | 2 | x |
| Poales | *Zea mays* | 2 | x |
| Poales | *Andropogon gerardi* | 3 | x |
| Poales | *Miscanthus sinensis* | 2 | x |
| Poales | *Panicum virgatum* | 2 | x |
| Poales | *Saccharum officinarum* | 6 | x |
| Caryophyllales | *Chenopodium quinoa* | 2 | x |
| Gentianales | *Coffea arabica geisha* | 2 | x |

**Table 2:(continued)**
| **Taxon** | **Species** | **#CAO genes** | **Tandem** |
| --- | --- | --- | --- |
| Apiales | *Hydrocotyle leucocephala* | 2 | x |
| Boraginales | *Ehretia anacua* | 2 | x |
| Asterales | *Helianthus annuus* | 2 | x |
| Lamiales | *Olea europaea* | 2 | x |
| Saxifragales | *Kalanchoe fedtschenkoi* | 2 | x |
| Saxifragales | *Kalanchoe laxiflora* | 4 | x |
| Fabales | *Arachis hypogaea* | 2 | x |
| Fabales | *Chamaecrista fasciculata* | 2 | x |
| Fabales | *Glycine max* | 4 | x |
| Fabales | *Glycine soja* | 4 | x |
| Fabales | *Lupinus albus* | 2 | x |
| Fabales | *Phaseolus vulgaris* | 2 | x |
| Fagales | *Carya illinoensis* | 2 | x |
| Rosales | *Fragaria x ananassa* | 3 | x |
| Rosales | *Malus domestica* | 2 | x |
| Malpighiales | *Linum usitatissimum* | 4 | x |
| Malpighiales | *Populus deltoides* | 2 | x |
| Malpighiales | *Populus nigra x maximowiczii* | 2 | x |
| Malpighiales | *Populus tremula x alba* | 2 | x |
| Malpighiales | *Populus trichocarpa* | 2 | x |
| Malpighiales | *Salix purpurea* | 2 | x |
| Sapindales | *Anacardium occidentale* | 3 | x |
| Malvales | *Gossypium barbadense* | 4 | x |
| Malvales | *Gossypium darwinii* | 4 | x |
| Malvales | *Gossypium hirsutum* | 4 | x |
| Malvales | *Gossypium mustelinum* | 4 | x |
| Malvales | *Gossypium raimondii* | 2 | x |
| Malvales | *Gossypium tomentosum* | 4 | x |
| Brassicales | *Alyssum linifolium* | 2 | x |
| Brassicales | *Caulanthus amplexicaulis* | 2 | x |
| Brassicales | *Iberis amara* | 2 | x |
| Brassicales | *Lepidium sativum* | 2 | x |
| Brassicales | *Stanleya pinnata* | 2 | x |
| Brassicales | *Brassica juncea* | 3 | x |
| Brassicales | *Brassica oleracea* | 2 | x |
| Brassicales | *Cakile maritima* | 2 | x |
| Brassicales | *Crambe hispanica* | 2 | x |
| Brassicales | *Eruca vesicaria* | 2 | x |
| Brassicales | *Sinapis alba* | 3 | x |
| Brassicales | *Camelina sativa* | 3 | x |
| Brassicales | *Isatis tinctoria* | 4 | x |

We also detected multiple occurrences of heterodimeric *CAO* genes, where the functional CAO protein is encoded by two genes (Table 1, Supplementary Table S1). Previous work in the prasinophyte *Micromonas pusila* demonstrated that in this species CAO was encoded by two separate genes, one that possesses the mononuclear iron-binding motif (*MpCAO1*) and a second one that encodes for the Rieske cluster (*MpCAO2*). It was shown that a combination of both gene products was necessary to form an active heterodimeric CAO protein (Kunugi et al. 2013; Dey et al. 2023). We expand on this work by showing the presence of a heterodimeric CAO encoded by two separate genes across the Mammiellophycea. This group within the prasinophytes contains *M. pusila* and several of its relatives (Worden et al. 2009), the ecologically important picoalgae *Ostreococcus tauri*, *Ostreococcus lucimarinus* and *Bathycoccus prasinos* (Palenik et al. 2007; Moreau et al. 2012; Blanc-Mathieu et al. 2014), as well as the related *Chloropicon primus* (Chloropicophyceae) (Lemieux et al. 2019).

We also show that heterodimeric CAO is not restricted only within the Prasinophytes. We detect a similar *CAO* gene arrangement in the genome of the multinucleate green alga *Caulerpa lentillifera* (Bryopsidales) (Arimoto et al. 2019), as well as in in the genome of *Bigelowiella natans* (Chlorarachniophyceae) (Curtis et al. 2012). Phylogenetic analysis based on plastid-encoded protein sequences revealed that chlorarachniophytes acquired their secondary plastids by the uptake of a filamentous green alga in the Bryopsidales (Suzuki et al. 2016). Thus, *B. natans* likely gained a heterodimeric CAO from the Bryopsidales endophyte. Considering that CAO in all other green lineages (Supplementary Table 1) and within the prochlorophytes (Nagata et al. 2004) is also encoded by a single gene that contains both the Rieske and iron-binding domain it is likely that the occurrence of two separate *CAO* genes is not an ancestral condition. Instead, it has been postulated that CAO was encoded by a single gene in the common ancestor of green plants (Kunugi et al. 2016). Indeed, it has been suggested that acquisition of two separate genes encoding for key functional domains may not be an evolutionarily difficult process. CAO likely functions as a trimer (Kunugi et al. 2013; Liu et al. 2022a), and the electron transfer from the Rieske to the mononuclear iron occurs between two neighbouring CAO proteins. A functionally similar process likely occurs in the CAO enzyme encoded by two separate genes.

All examined streptophytes encode for at least one *CAO* gene (Supplementary Table 1) signifying the importance of Chl *b* for plant physiology, but we were not able to identify a homolog of *CAO* in several chlorophyte algal species. Not surprisingly, we did not detect a *CAO* homolog in the invertebrate parasite *Helicosporidium* sp., a colorless protist that belongs within the trebouxiophycean green algal clade (Tartar et al. 2003). As a consequence of its parasitic lifestyle, *Helicosporidium* has lost nearly all genes associated with light harvesting, photosynthesis and chlorophyll biogenesis (Pombert et al. 2014), including CAO.

We were also not able to conclusively identify a *CAO* gene in the genomes of the endolithic coral holobiont *Ostreobium quekettii* (Bryopsidales) (Iha et al. 2021), the aquaculture species *Tetraselmis striata* (Chlorodendrophyceae) (Steadman Tyler et al. 2019) and the halotolerant *Picocystis* sp. ML (Picocystophyceae) (Junkins et al. 2019). All identified sequences within the genomes of these species had e-values higher than the cut-off (≥e-15) used in this work and high similarity to pheophorbide *a* oxygenase (PAO), a Rieske-type protein involved in chlorophyll degradation (Pružinská et al. 2003). The lack of *CAO* genes in these algae is surprising and could be an artifact of genome assembly, as Chl *b* has been detected in all three species (Roesler et al. 2002; Massé et al. 2020; Conlon et al. 2024). The genome of *O. queketti* has a low BUSCO score (60.7%) (Iha et al. 2021) and the BUSCO score is not reported for the draft genomes of *T. striata* and *P.* sp. ML (Junkins et al. 2019; Steadman Tyler et al. 2019). The presence of CAO should be experimentally examined in these species.

It is tempting to speculate that these species may have a different mechanism for Chl *b* biosynthesis or a CAO sequence significantly different than most algae and plants. For instance, *Ostreobium* is an endolithic coral holobiont adapted to extreme shading, and has lost many genes associated with photoprotection and light perception due to evolution in a light-limited environment (Iha et al. 2021). This alga is also the only known eukaryote that has lost the gene encoding light-dependent protochlorophyllide oxidoreductase (LPOR), a key enzyme that catalyzes the conversion of protochlorophyllide to chlorophyllide *a*, a necessary step towards the synthesis of Chl *a* (Reinbothe et al. 2010). Instead, this alga fully depends on dark operative POR (DPOR) to synthesize Chl *a*, likely as an adaptation to extreme shading (Iha et al. 2021). The mechanism of chlorophyll biosynthesis has not been examined in the species in detail, but it is clear that the endolithic lifestyle has resulted in a unique physiology. Further work will shed light on the role of CAO in this and other shade-tolerant species.

We used a very conservative method to identify putative CAO sequences with both a Rieske and Fe-binding domains, but this strict approach may have prevented the identification of duplicates that may have diverged or become pseudogenes. Indeed, using a less conservative cutoff (≤e-5) when screening the genomes of *A. thaliana* and *C. reinhardtii* (Wang et al. 2022; Craig et al. 2022) reveals that *PAO* and Translocon at the inner envelope membrane of chloroplasts 55 (*TIC55*) may be paralogous to *CAO* (Supplementary Table S2). *PAO* and *TIC55* encode for enzymes with a Rieske and Fe-binding domains but share a very low sequence similarity with CAO and each other (Supplementary Table S2, Supplementary Figure S1). Both PAO and TIC55 are involved in chlorophyll breakdown and allow for the degradation of the highly phototoxic pheophorbide *a* and its export from the chloroplast (recently reviewed in Kuai et al. 2018). Evolutionary analysis of the genes involved in chlorophyll degradation revealed that this pathway was already present in the common ancestor of land plants, suggesting that *PAO* and *TIC55* may share their origins with *CAO* (Schumacher et al. 2022). This hypothesis has not yet been experimentally supported as ancestral monooxygenase have been difficult to distinguish without detailed knowledge on their biochemical activity and function (Schumacher et al. 2022). This possibility raises interesting questions about the ancient evolutionary origins of chlorophyll metabolism within the Viridiplantae.

### Was CAO gene duplication an ancestral event?

To determine whether the *CAO* gene duplication was an ancestral event, we performed a phylogenetic analysis of all high-quality CAO amino acid sequences (except heterodimeric CAO). Phylogenetic patterns typically match previously shown relationships among major Viridiplantae groups (Figure 2), although some relationships are poorly resolved (particularly among the algal groups and the Rosids; Supplementary Figure 2). The presence of two *CAO* genes has been previously reported in rice (*OsCAO1* and *OsCAO2*) where the two genes were positioned in tandem on Chromosome 10, taken as evidence for a recent gene duplication in this species (Lee et al. 2005). A tandem CAO duplication, however, appears to be the exception rather than the rule. In addition to rice, we only observed *CAO* duplicates located in proximity to each other in the genomes of a handful of species: the green alga *Volvox reticuliferus*, the lycophyte *Diphastrium complanatum*, and the magnoliid *Cinnamomum kanehirae* (Table 2).

**Figure 2:**
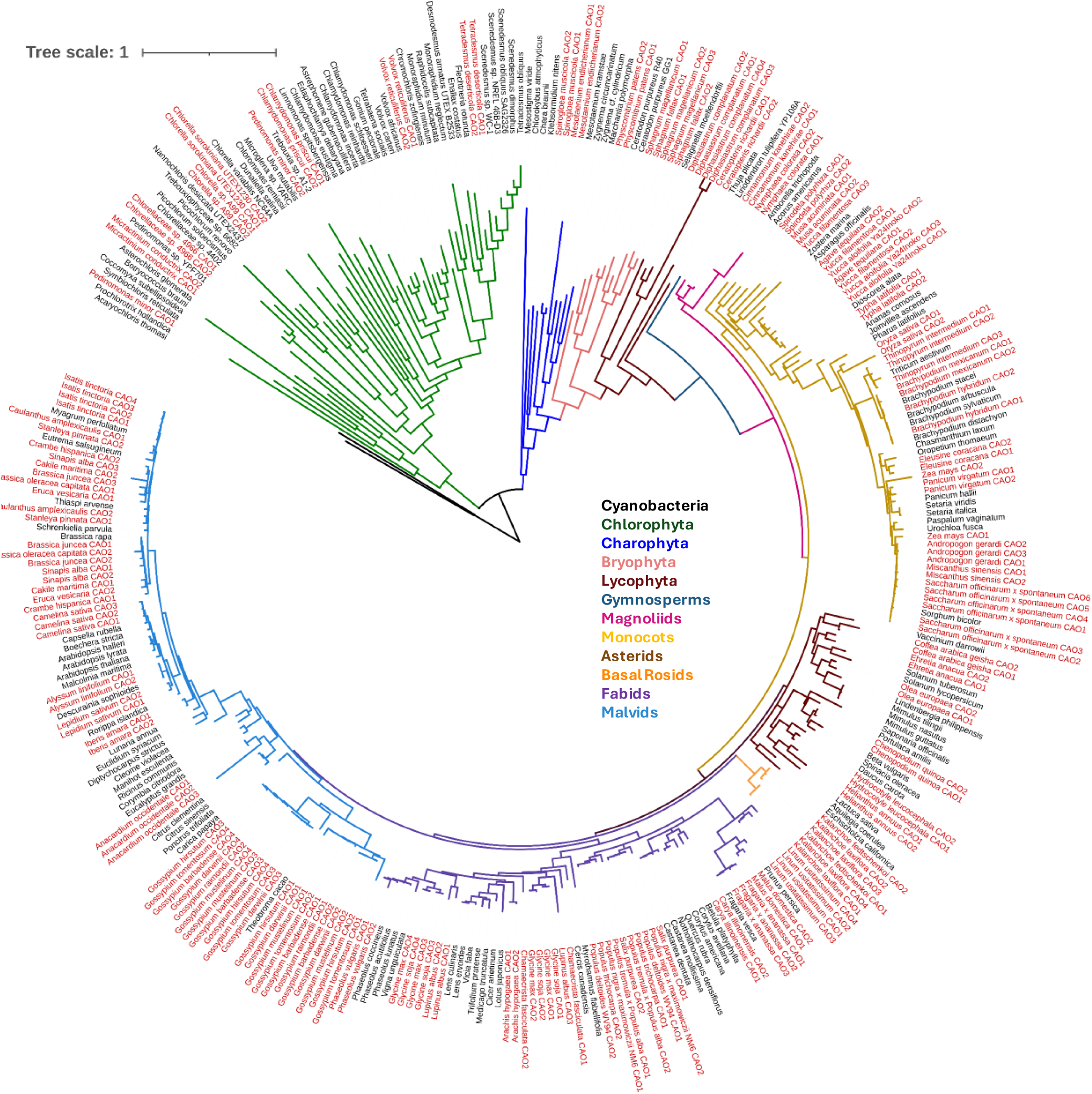
A phylogenetic tree based on CAO amino acid sequences inferred from using maximum likelihood analysis, using 326 unique CAO sequences across 246 species color-coded to represent the major Viridiplantae groups. Scale bar indicates amino acid substitutions per site. Species with more than one detected CAO gene are highlighted in red

There are several instances, mainly in very closely related species, where the duplication may have originated in a recent shared ancestor. There are several possible cases in the Trebouxiophyceae and Chlorophyceae class, where *CAO* duplicates are detected in closely related *Chlorella* and *Scenedesmus* species (Supplementary Figure 2A). This insight must be interpreted with caution as phylogenetic resolution within the chlorophytes is still weak due to limited gene and taxon sampling (Li et al. 2021). Within the bryophytes, *Spaghnum magellanicum* and *Sphagnum falax*, species that share a very close evolutionary relationship (Piatkowski and Shaw 2019; Bell et al. 2020), both have two *CAO* genes possibly arising from an ancestral duplication. Two CAO genes are also encoded in the genome of *P. patens* (Zhang et al. 2023). This raises the possibility of a CAO duplication in a recent shared ancestor (Supplementary Figure 2B), but the bryophytes are represented by only 5 species in our analysis. Additional genomic sequencing and a deeper phylogenetic analysis will be needed to clarify the evolution of CAO within these groups.

Species that encode for multiple *CAO* genes within their genomes are more prevalent among angiosperms. Within the monocots, *CAO* duplicates arising from a shared ancestral gene are likely present in the Asparagaceae, specifically among the closely related *Agave tequiliana*, *Yucca filamentosa* and *Yucca aloifolia* (Ji et al. 2023), although we were only able to detect a single *CAO* copy in *Asparagus officinalis* (Supplementary Figure 2C). We detected two *CAO* gene copies in two *Brachypodium* species (*B. mexicanum* and *B. hybridum*) but not in their relatives from the same genus (*B. arbuscula*, *B. stacei*, *B. sylvaticum* and *B. distachyon*), raising the possibility of an ancestral duplication followed by gene loss. All phylogenetic studies in *Brachypodium* support a rapid and recent divergence from a common ancestor (Catalan et al. 2016), supporting this possibility.

*CAO* duplication may be ancestral in the two closely related Saxifragales *Kalanchoe laxiflora* and *Kalanchoe fedtschenkoi* (Han et al. 2024). We detected two *CAO* genes in *K. fedtschenkoi* and four in *K. laxiflora*, possibly arising in a recent ancestor, followed by a second duplication within *K. laxiflora* (Supplementary Figure 2D). Within the Fabids, four *Populus* species and their relative *Salix purpurea*, each encode for two *CAO* gene copies that may be a result of an ancestral duplication. The cultivated soybean *Glycine max* and its wild relative *Glycine soja* each encode for four *CAO* gene copies (Supplementary Figure 2D) that are closely related to their homologs between the two species. The phylogenetic separation of *CAO1/2* and *CAO3/4* from *G. max* and *G. soja* is puzzling, as the two species are the closest relatives within the *Glycine* genus (Zhuang et al. 2022). All *Gossypium* species examined here encode for four *CAO* genes, except *G. raimondii* which encodes for two copies (Supplementary Figure 2E). The only other member of the Malvaceae examined, *Theobroma cacao,* encodes for a single *CAO* gene, and although it clusters with *CAO* from *Gossypium* sp., the relationship is poorly supported. Finally, there is a possibility of ancestral duplications within the Brassicaceae (Supplementary Figure 2E), but this family has a poor resolution due to factors such as incomplete sampling, rapid radiation, and hybridization (Huang et al. 2016; Hendriks et al. 2023).

The genomes of multiple species across the Viridiplantae encode for more than one *CAO* gene, but are these gene duplications related to plant evolutionary patterns? The green lineage is characterized by a dynamic and complex evolutionary history marked by multiple whole genome (WGD) and segmental duplication events, gene family expansions, and lineage-specific gene losses (Panchy et al. 2016; Kersey 2019; Leebens-Mack et al. 2019; Soltis and Soltis 2021). Such tremendous diversity in genome content can be a challenge for elucidating *CAO* evolutionary trajectories even withing a single family. For instance, the Brassicaceae have been shaped by multiple well-established WGD events (Mabry et al. 2020), but we detected only a single *CAO* gene in the genome of *A. thaliana* and its closest relatives. In contrast, the genomes of several other Brassica species encode for two or more CAO genes (Figure 2). This highlights the importance of considering not only gene duplication but also gene retention dynamics when interpreting the evolution of CAO. Future work that incorporates in-depth analyses, such as relative dating of CAO duplication events and genome-wide synteny, will further clarify the evolutionary trajectory of CAO within the Viridiplantae.

We show that *CAO* gene duplication is not a rarity among the Viridiplantae, but are all *CAO* duplicates expressed and do they encode for active proteins? In our work, we focused only on CAO genes with highly conserved catalytic domains likely to encode for active enzymes, but the functional role of CAO duplication may be complex. A survey of published transcriptomes from 16 representative species from all major groups (Chlorophytes, Charophytes, non-vascular and vascular plants) suggests that all *CAO* duplicate genes within a species are expressed, albeit at different levels. Typically, one *CAO* gene has a higher expression than its paralog within the same genome (Supplementary Table 4) suggesting differential regulation of *CAO* expression.

Significantly, all experimental studies to date suggest that the expression of *CAO* duplicates may be differently regulated by light availability. In rice, the expression of *OsCAO1* is light-inducible, while *OsCAO2* accumulates during periods of darkness and is expressed mainly in non-photosynthetic tissues (Lee et al. 2005). In *P. patens*, both *PpCAO1* and *PpCAO2* are expressed in both light and dark conditions, but only *PpCAO1* is upregulated with prolonged light exposure.

Similarly, in two *Chlamydomonas* species that harbour *CAO* duplicate genes (*Chlamydomonas priscui* and *Chlamydomonas* sp. ICE-MDV), only one *CAO* homolog is upregulated in response to increases in light intensity, while the second one is constitutively expressed (Poirier et al. 2025). The role of light in regulating *CAO* expression has been also demonstrated previously in *Arabidopsis* and tobacco, species with a single CAO gene copy (Pattanayak et al. 2005; Tanaka and Tanaka 2005; Biswal et al. 2012, 2024).

Additionally, the mechanisms behind the control of Chl *b* synthesis may be species specific. Mutational analysis in rice revealed that only *OsCAO1* was necessary for Chl *b* biosynthesis. *OsCAO1* knockout mutants exhibited low Chl *b* levels, disordered thylakoid membrane arrangements, membrane damage due to excessive ROS accumulation and decreased chilling tolerance (Lee et al. 2005; Jung et al. 2021; Xiong et al. 2024), but these effects were absent in *OsCAO2* knockouts (Lee et al. 2005). In contrast, knocking out either of the two *CAO* homologs in *P. patens* lead to significantly reduced Chl *b* levels suggesting a dependency on two active CAO enzymes for the appropriate control of Chl *b* synthesis in this species (Zhang et al. 2023).

It is tempting to speculate that species that encode for multiple *CAO* genes have a more robust capacity for Chl *b* biosynthesis through an increase in gene dosage or a finer control over Chl *b* accumulation though gene subfunctionalization or differential expression (Kondrashov 2012; Panchy et al. 2016). Two gene copies with different expression patterns or activity levels could provide a fine-tuned control of Chl *b* levels and photosynthetic performance under varying environmental conditions, but additional experimental work is needed to substantiate this hypothesis.

### Evolutionary implications of the CAO degron sequence

In addition to light-dependant regulation at the level of transcription, the main regulation of CAO occurs post-translationally. Early work in *A. thaliana* demonstrated that this regulation is governed by the presence of a degron sequence (QDLLTIMILH) in the protein’s N-terminal A domain that acts as a docking site for a chloroplast Clp protease (Yamasato et al. 2005; Nakagawara et al. 2007; Sakuraba et al. 2009). It has been hypothesized that the degron sequence is “hidden” within the A domain when there are insufficient amounts of Chl *b*, but accumulation of this product causes a conformational change in the CAO protein, exposing the degron, and giving access to the protease (Tanaka and Tanaka 2011). The evolution of this negative-feedback regulatory loop has not been examined in detail, but it has been suggested that the degron was acquired early in the Viridiplantae evolution as a mechanism for high light adaptation (Kunugi et al. 2016).

Here we show that the degron sequence is highly conserved among all land plants (86% identity), moderately conserved among charophytes and non-vascular plants (68% and 66% identity respectively), and very poorly conserved in chlorophytes (39% identity) (Figure 3; Supplementary Figure 3). Land plants (both vascular and non-vascular) have several highly conserved sites across all species (Asparagine at position 2, Isoleucine at position 6, Histidine at position 10), whereas these positions are only moderately conserved in charophytes, and poorly conserved in chlorophytes. In contrast, the Rieske cluster [CXH(X)15-17CXXH] exhibits an 84% pairwise identity among all species, where the two histidine (positions 3 and 23) and cysteines (positions 1 and 20) that coordinate the metallocentre [2Fe-2S] (Przybyla-Toscano et al. 2021) are present in all examined sequences. Nearly all residues are conserved among the Viridiplantae in the C-terminal Fe-binding domain (96% pairwise identity). Overall, these results suggest that the mechanism of Chl *b* biosynthesis involving the Rieske and Fe-binding domains is ancestral and conserved among all photosynthetic species, but this is not the case for the regulatory degron domain.

**Figure 3:**
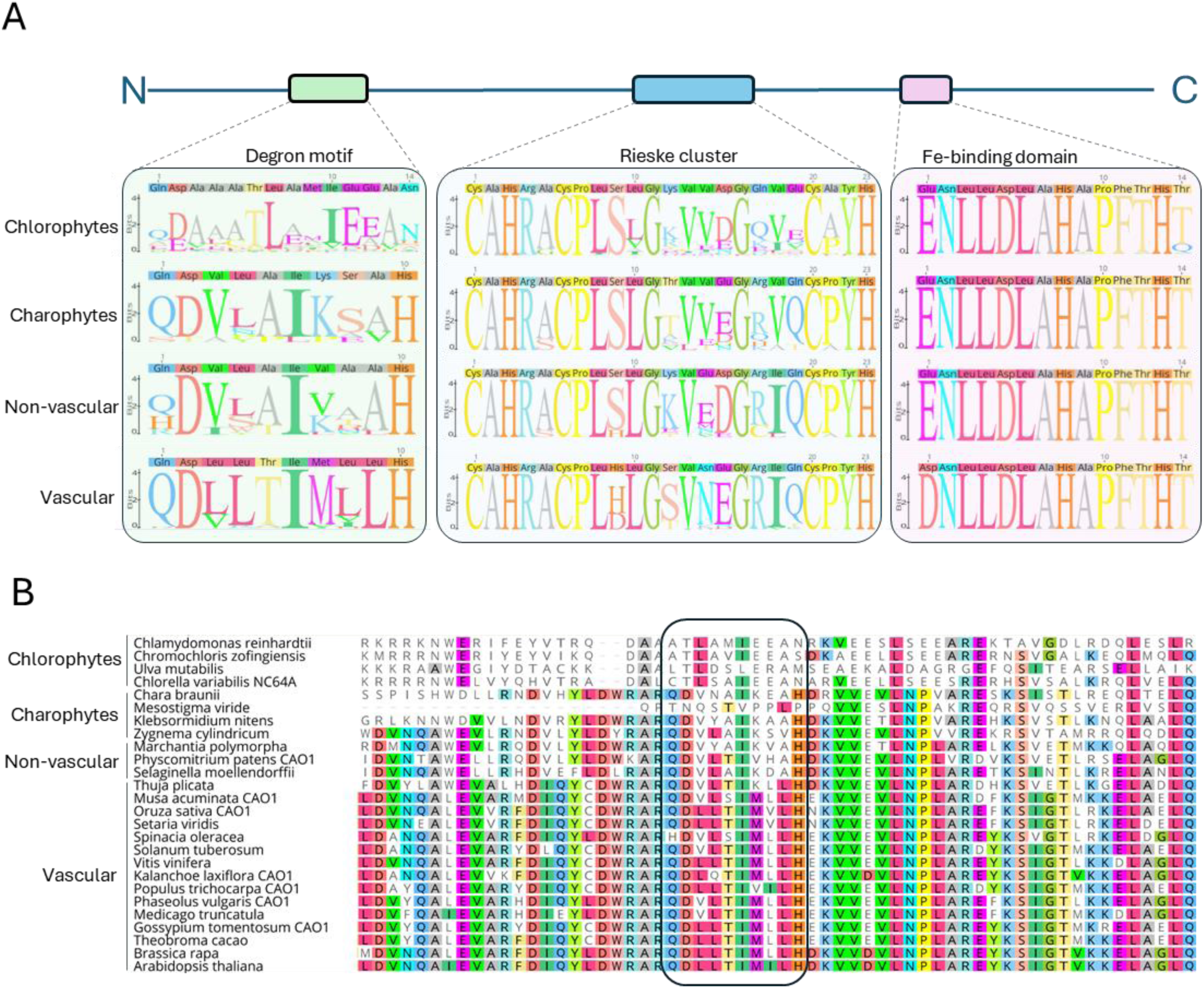
Key domains in the CAO protein across the Viridiplantae. (A) Sequence logos of the conserved motifs identified in the CAO protein sequences. N and C represent the N-terminus and C-terminus, respectively. All sequence logos are superimposed on a graphical representation of plant CAO sequences (not to scale) showing the regulatory degron motif, conserved Rieske domain cluster and Fe-binding domain (Sakuraba et al., 2009; Liu et al., 2022). The logos were generated for CAO proteins from chlorophyte algae, charophyte algae, non-vascular plants, and vascular plants. The sequences on top of each logo are the consensus sequence with the most common base in each sequence alignment. (B) Comparison of the CAO amino acid sequences corresponding to the degron and its surrounding region in the N-terminal regulatory domain in representative species from Chlorophyte, Charophyte, non-vascular and vascular land plants. The region that corresponds to the degron sequence as described in *A. thaliana* is emphasized with a box. Amino acids that are identical with the *A. thalina* sequence are colored, and those that are different are shown in white.

The lack of a highly conserved degron sequence may have several implications in the evolution of efficient light harvesting. For instance, the N-regulatory domain (including the degron) is completely absent in marine eukaryotic Mammiellophycea that encode for a heterodimeric *CAO* gene, as well as in their deep-water relatives from the order Palmophyllales. These species typically have low Chl *a/b* ratios and, unlike land plants, can incorporate Chl *b* into their core antennae. This was interpreted as a necessary adaptation for efficient light harvesting in deep marine environments characterized by low levels of blue-green light (Kunugi et al. 2016).

It has been suggested that the degron sequence evolved in shallow-water algae and land plants as a means to regulate Chl *b* accumulation and limit Chl *b* incorporation into the reaction center in an environment characterized with high light intensities (Kunugi et al. 2016). Indeed, such species exclusively harbour Chl *a* in their core complexes, but this appears to be controlled by pigment availability rather than the chemical nature of the two chlorophyll types. The binding affinity of Chl *a* and *b* to the PSI complex is similar (Ikegami et al. 2007) and cyanobacteria that do not naturally produce Chl *b* successfully incorporated this pigment into their reaction centers when transformed with plant CAO (Satoh et al. 2001; Xu et al. 2001). Incorporation of Chl *b* into the reaction center accompanied by increased Chl *b* accumulation was also observed in *Arabidopsis* when full length *CAO* was overexpressed or replaced with a prokaryotic *CAO* that lacked a degron sequence (Hirashima et al. 2006; Sakuraba et al. 2009).

The chlorophyll cycle has not been examined in detail in the core chlorophytes (Chlorophyceae, Trebouxiophyceae, Ulvophyceae), a diverse group that inhabits marine, freshwater and terrestrial habitats (Leliaert et al. 2012). Although many chlorophytes thrive in high light, others are adapted to light limited environments, including deep waters. As a result, these species have developed adaptations allowing for efficient light harvesting in low light. For instance, *Chlamydomonas priscui* is endemic to the depths of the Antarctic Lake Bonney, an environment characterized by extreme shading and dominated by blue-green wavelengths (Neale and Priscu 1995). This alga displays a host of adaptations to extreme shading including a large LHC antennae size (reviewed in Cvetkovska et al. 2017; Hüner et al. 2022). As suggested for marine prasinophytes (Kunugi et al. 2016), such low-light adapted chlorophytes could benefit from increased Chl *b* levels and could thrive without a Chl *b*-dependent feedback mechanism.

Despite the lack of a conserved degron sequence to regulate CAO protein levels (Figure 3), chlorophytes have maintained the ability to alter their Chl *a/b* ratio in response to light (Leliaert et al. 2012). Transcriptional control of CAO levels was postulated to be a minor aspect of adjusting Chl *b* levels in plants (Tanaka and Tanaka 2019), but this process could play a bigger role in chlorophytes. It is also possible that a different, yet unidentified, mechanism exists for Chl *b* turnover independent of a degron sequence, allowing these species to modify their Chl *a/b* ratios. Finally, we can not rule out that the region corresponding to a degron in Chlorophyte CAO may still have a similar function despite poor sequence conservation with land plants. The mechanism of regulating protein degradation via degron motifs is well understood in plants (Isono et al. 2024), but insights on algal degrons is lacking. A detailed experimental investigation on the control and mechanism of Chl *b* biosynthesis is clearly needed among the Chlorophytes. For instance, targeted mutagenesis of the N-terminus of algal CAO will reveal the importance and role of this regulatory region in chlorophyll turnover.

### Conclusion and Drawbacks

Our work suggests that *CAO* gene duplication is widespread among the Viridiplantae, likely originating from multiple independent duplication events throughout plant evolution. While we show that the presence of multiple *CAO* gene copies is highly prevalent in land plants, these results must be interpreted with caution. Many more angiosperm genomes are available in public databases, while other plant and algal groups are not as well represented. Thus, the conclusions of our study must be re-visited once more genomes are sequenced. Furthermore, many genome sequencing initiatives are focused on plants of economic and agricultural importance, in which domestication and selective breeding may have affected the genomic content (Turner-Hissong et al. 2020). While the exact physiological impact of CAO duplication is unknown, a study in transgenic tobacco overexpressing CAO (Biswal et al. 2012) showed that an increase in Chl *b* may be a strategy to enhance CO2 assimilation, delay senescence, and enhance productivity. Such traits would be attractive in crop breeding programs (Voitsekhovskaja and Tyutereva 2015), and the high prevalence of CAO duplication seen in crop plants in this study may have been selected for during plant breeding. Indeed, several crop plants (including corn, soybean, and olives, among others) appear to express all CAO genes encoded in their genome (Supplementary Table 4). The functional role of these duplicates in plant physiology, photosynthetic efficiency, and productivity warrants further investigation.

Our understanding of the prevalence and evolution of CAO at the genetic level will be improved in future years, as more algal and non-vascular plant genomes become available. Furthermore, there are still numerous complex and outstanding questions regarding the control of Chl *b* accumulation in photosynthetic species. In particular, the chlorophytes are an understudied group when it comes to Chl *b* regulating mechanism. Detailed genomic and experimental studies will shed light on the role and regulation of Chl *b* in these species and will deepen our understanding on this crucial photosynthetic pigment.

## Acknowledgements

This project was supported by Natural Sciences and Engineering Research Council of Canada Discovery Grants (NSERC DG) awarded to M.C. The authors are grateful for the support from the Canada Foundation for Innovation (CFI) and University of Ottawa start-up funding. M.P. was supported by Ontario Graduate Scholarship (OGS), NSERC Graduate Scholarship, and Polar Knowledge Canada Antarctic Doctoral Scholarship.

## Author’s contributions

M. Poirier and M. Cvetkovska conceptualized the work and designed the experiments. Material preparation, data collection and analysis were performed by M. Poirier, R. Wright and M. Cvetkovska. The first draft of the manuscript was written by M. Poirier and all authors commented on all versions of the manuscript. All authors read and approved the final manuscript.

## Supplementary Information

**Supplementary Table S1**: A summary of genes encoding for chlorophyllide *a* oxygenase (*CAO*) across the Viridiplantae used in this work. All sequences were obtained from Phycocosm (Chlorophytes, Euglenophytes, Cercozoa, Charophytes) or Phytozome v13 (Bryophytes, Marchantiophytes, Lycophytes, Gymnosperms, Angiosperms). Accession numbers represent the unique IDs assigned to each gene (Phycocosm: Gene portal|Protein ID|Gene Model; Phytozome: Gene Model). E-values are based on a tBLASTn search with the *Chlamydomonas reinhardtii* CAO (Cre01.g043350) for species within the Chlorophyta and Charophyta, and the *Arabidopsis thaliana* CAO (AT1G44446) for all other species

**Supplementary Table S2:** BLAST results obtained by screening the genome of *Arabidopsis thaliana* and *Chlamydomonas reinhardtii* for CAO homologs. Accession numbers were obtained from Phytozome v13 (Goodstein et al. 2012). The functional domains according to InterPro (Blum et al. 2025) that are shared among all proteins are highlighted with an asterisk (Rieske Domain; InterPro: IPR017941; Pheophorbide a Oxygenase Domain; InterPro: IPR013626).

**Supplementary Table S3**: A summary of the occurrence of *CAO* duplicate genes across the major groups within Viridiplantae. The number in brackets represents the proportion of species which encode for multiple copies of the *CAO* gene (Mutiple CAO) or encode for two *CAO* genes that form heterodimeric complex (Heterodimeric CAO).

**Supplementary Table S4**: *CAO* gene expression in species whose genome encodes for more than one *CAO* gene. All data was obtained from previously published work. Gene expression is presented as FPKM, RPKM or TPM averages with standard deviation. In all cases, gene expression ratios were determined by dividing all expression values by the highest CAO expression level within the species.

**Supplementary Figure 1:** An alignment of the CAO protein from (A) *Arabidopsis thaliana* and (B) *Chlamydomonas reinhardtii* with all other proteins identified using the CAO amino acid sequence as a query. The color of the residue depends on the degree of conservation among proteins (blue fully conserved; green 80–100% similar; yellow 60–80% similar; white < 60% similar). The important domains that are shared among all proteins are highlighted in green (Rieske Domain; InterPro: IPR017941), yellow (Fe-binding domain) and blue (Pheophorbide a Oxygenase Domain; InterPro: IPR013626).

**Supplementary Figure 2:** A phylogenetic tree of CAO genes inferred from using maximum likelihood analysis. The node labels show the bootstrap support, with dashed lines indicating <50 bootstrap support. Scale bar indicates amino acid substitutions per site. Species with more than one detected CAO gene are highlighted in red. Cases where CAO duplication may have arisen in a common shared ancestor are marked with a blue arrow, and only those located on branches with a strong bootstrap support (>85%) are shown. The inset shows the outline of the full CAO phylogenetic tree inferred from using maximum likelihood analysis, using 326 unique CAO sequences across 246 species color coded to represent the major Viridiplantae groups. The position of the chlorophytes (green) is highlighted with a black box. (**A)** Chlorophytes; (**B**) Charophytes, bryophytes (including marachantiophytes), and lycophytes; (**C**) Gymnosperms, magnoliids, and monocots; (**D**) Basal eudicots, asterids and fabids; (**F**) Malvids.

**Supplementary Figure 3:** Multiple sequence alignment of the predicted CAO amino acid sequences from Chlorophytes (**A**), Charophytes (**B**), non-vascular plants (**C**) and vascular plants (**D**). Global alignments were performed with MUSCLE. The colour of the residue depends on the degree of conservation among species (blue, fully conserved; green, 80–100% similar; yellow, 60– 80% similar; gray, < 60% similar). As a reference, the CAO sequence from *Arabidopsis thaliana* is present in all alignments (top), with the key domains annotated (green, Degron sequence; blue, conserved Rieske cluster; purple, Fe-binding domain)

