## Supplementary Figure for "Chlorophyllide *a* oxygenase (CAO) gene duplication across the Viridiplantae"

### Supplementary data

**Mackenzie C. Poirier[1], Roberta Wright[1], Marina Cvetkovska[1]***

[1]Department of Biology, University of Ottawa, Ottawa, ON, Canada

***Corresponding author:**

**Marina Cvetkovska**

Department of Biology, University of Ottawa, 30 Marie-Curie Pr., Ottawa, ON, Canada, K1N 6N5

**ORCID:** 0000-0002-7080-8203

A

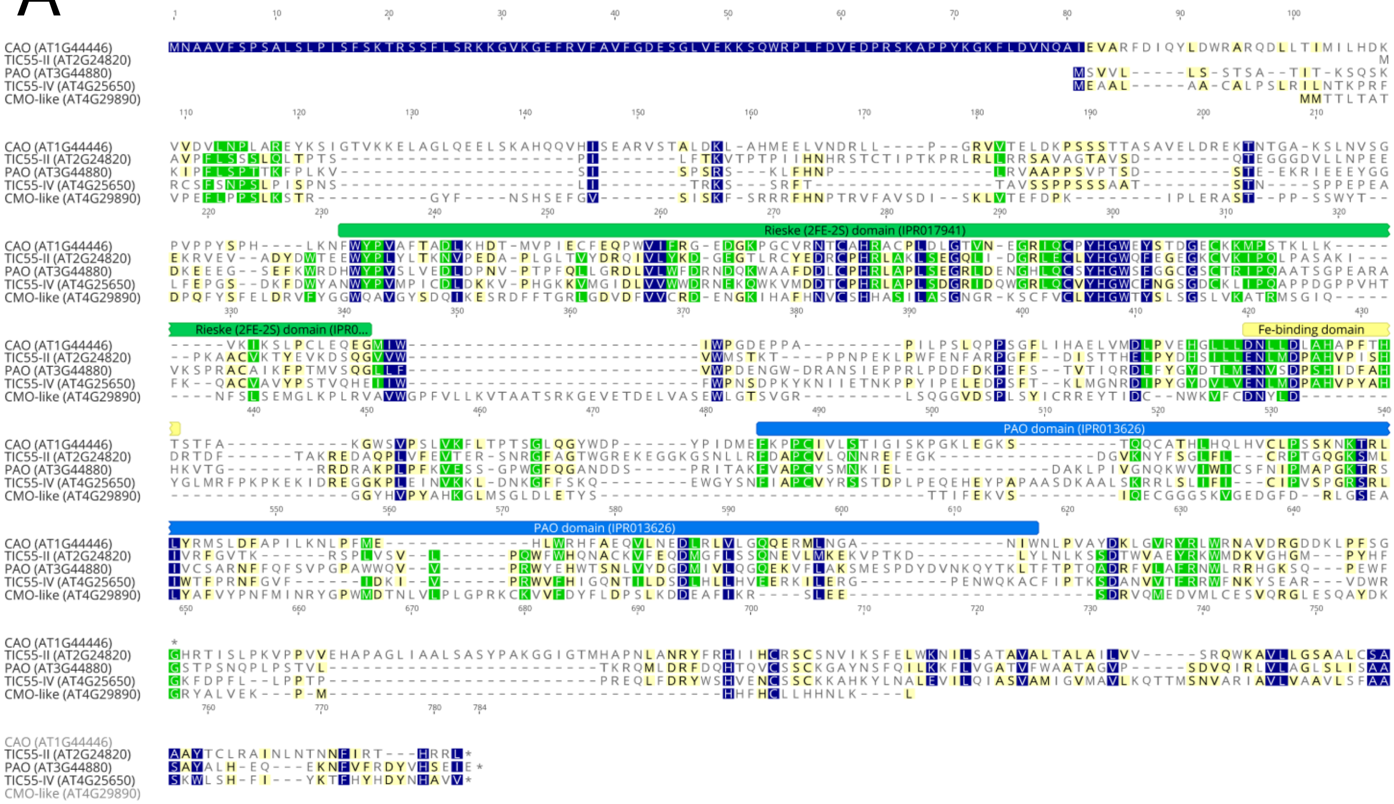

**Supplementary Figure 1**: An alignment of the CAO protein from (A) *Arabidopsis thaliana* and (B) *Chlamydomonas reinhardtii* with all other proteins identified using the CAO amino acid sequence as a query. The color of the residue depends on the degree of conservation among proteins (blue, fully conserved; green, 80–100% similar; yellow, 60–80% similar; white,< 60% similar). The important domains that are shared among all proteins are highlighted in green (Rieske Domain; InterPro:IPR017941), yellow (Fe-binding domain) and blue (Pheophorbide a Oxygenase Domain; InterPro:IPR013626) (Continued on the next page)

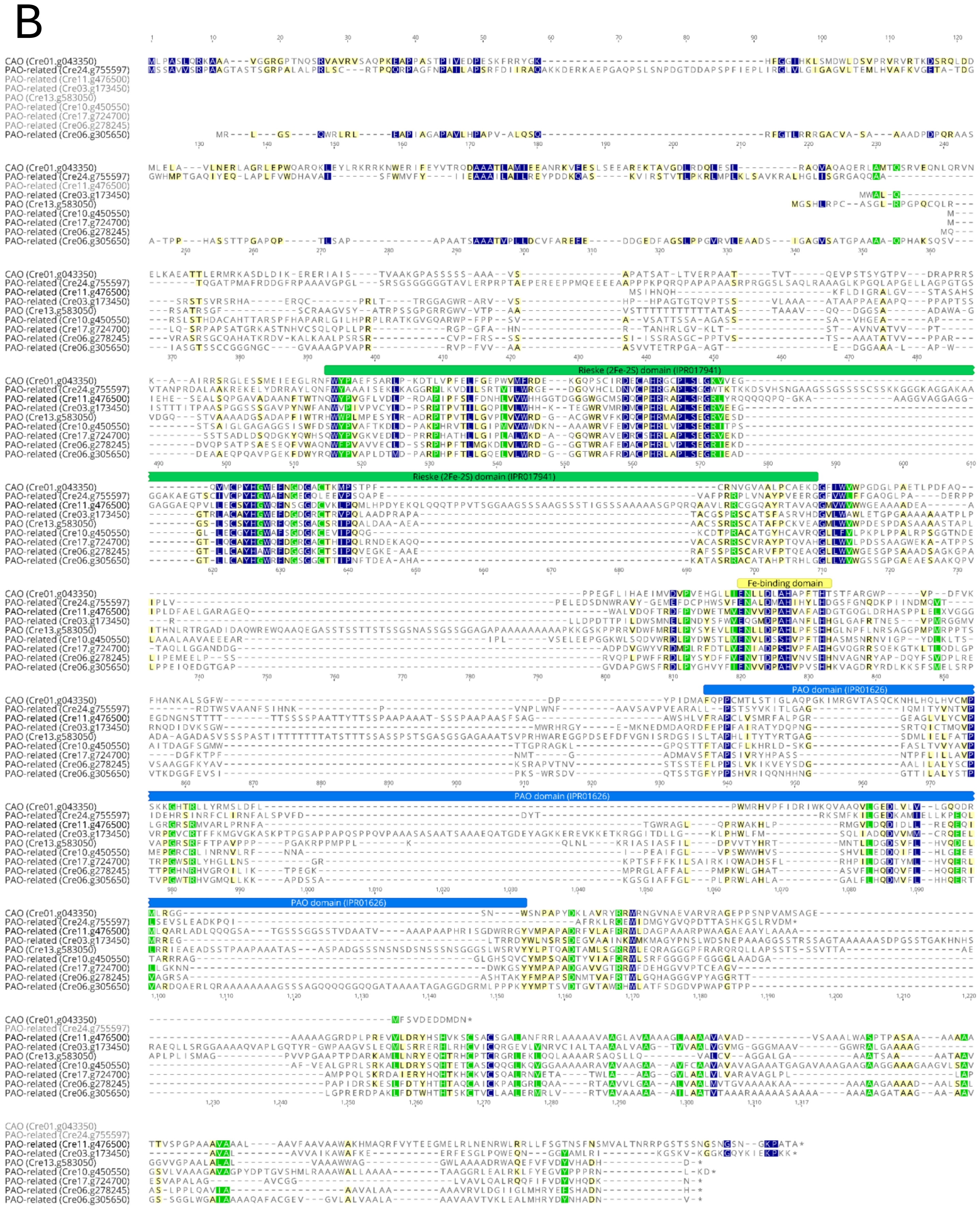


**Supplementary Figure 1:** An alignment of the CAO protein from ( (B) *Chlamydomonas reinhardtii*

# A

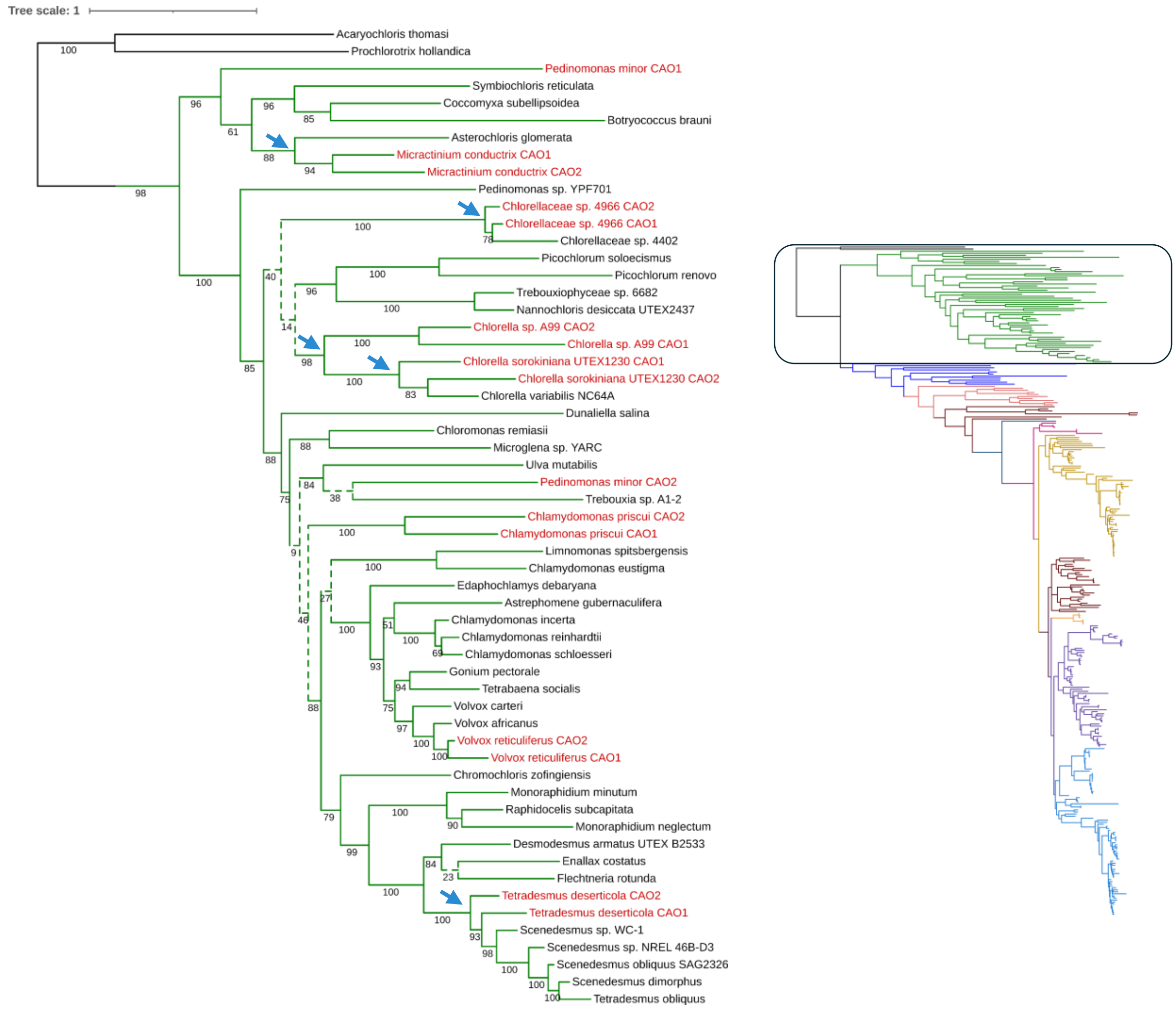


**Supplementary Figure 2**: A phylogenetic tree of CAO genes inferred from using maximum likelihood analysis. The node labels show the bootstrap support, with dashed lines indicating <50 bootstrap support. Scale bar indicates amino acid substitutions per site. Species with more than one detected CAO gene are highlighted in red. Cases where CAO duplication may have arisen in a common shared ancestor are marked with a blue arrow, and only those located on branches with a strong bootstrap support (>85%) are shown. The inset shows the outline of the full CAO phylogenetic tree inferred from using maximum likelihood analysis, using 326 unique CAO sequences across 246 species color coded to represent the major Viridiplantae groups. The position of the group is highlighted with a black box. (**A)** Chlorophytes

B

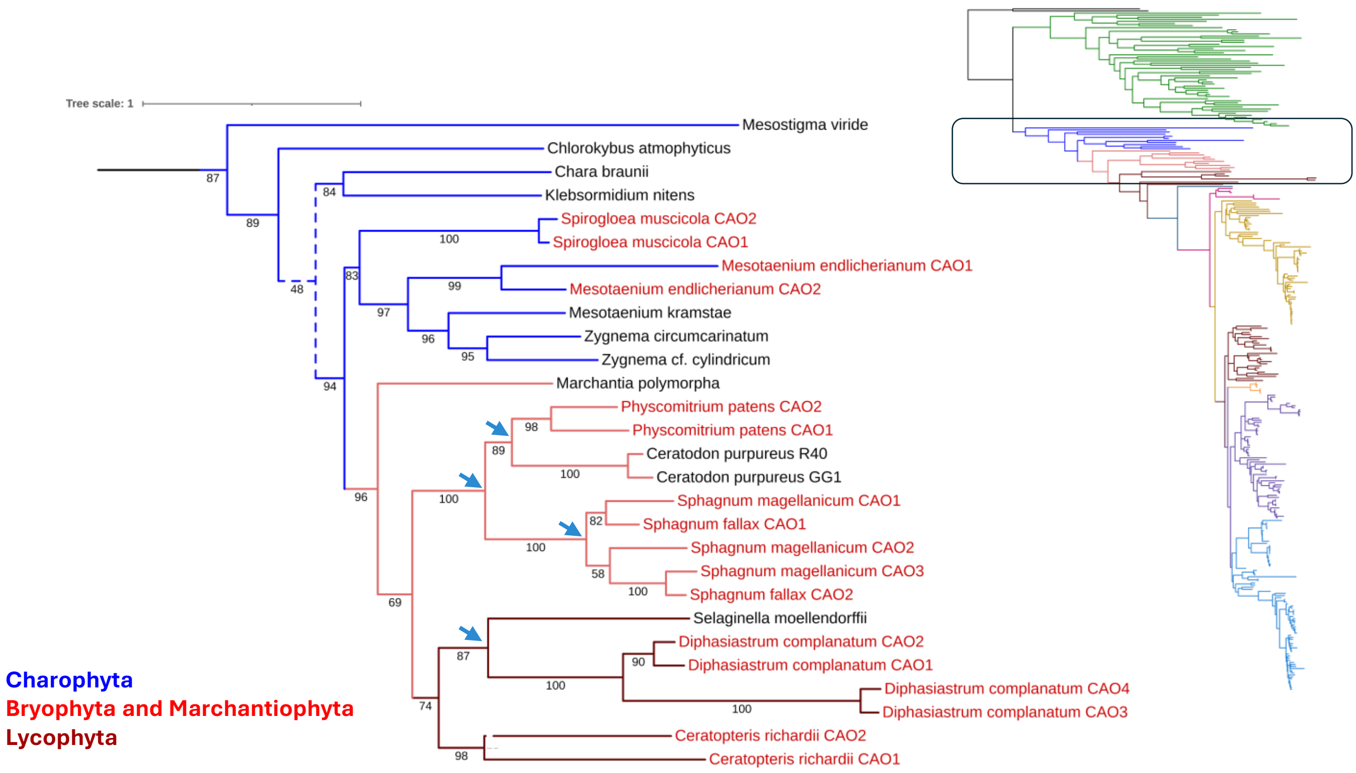


**Supplementary Figure 2**: (B) Charophytes, bryophytes (including marachantiophytes) and lycophytes. (Continued)

C

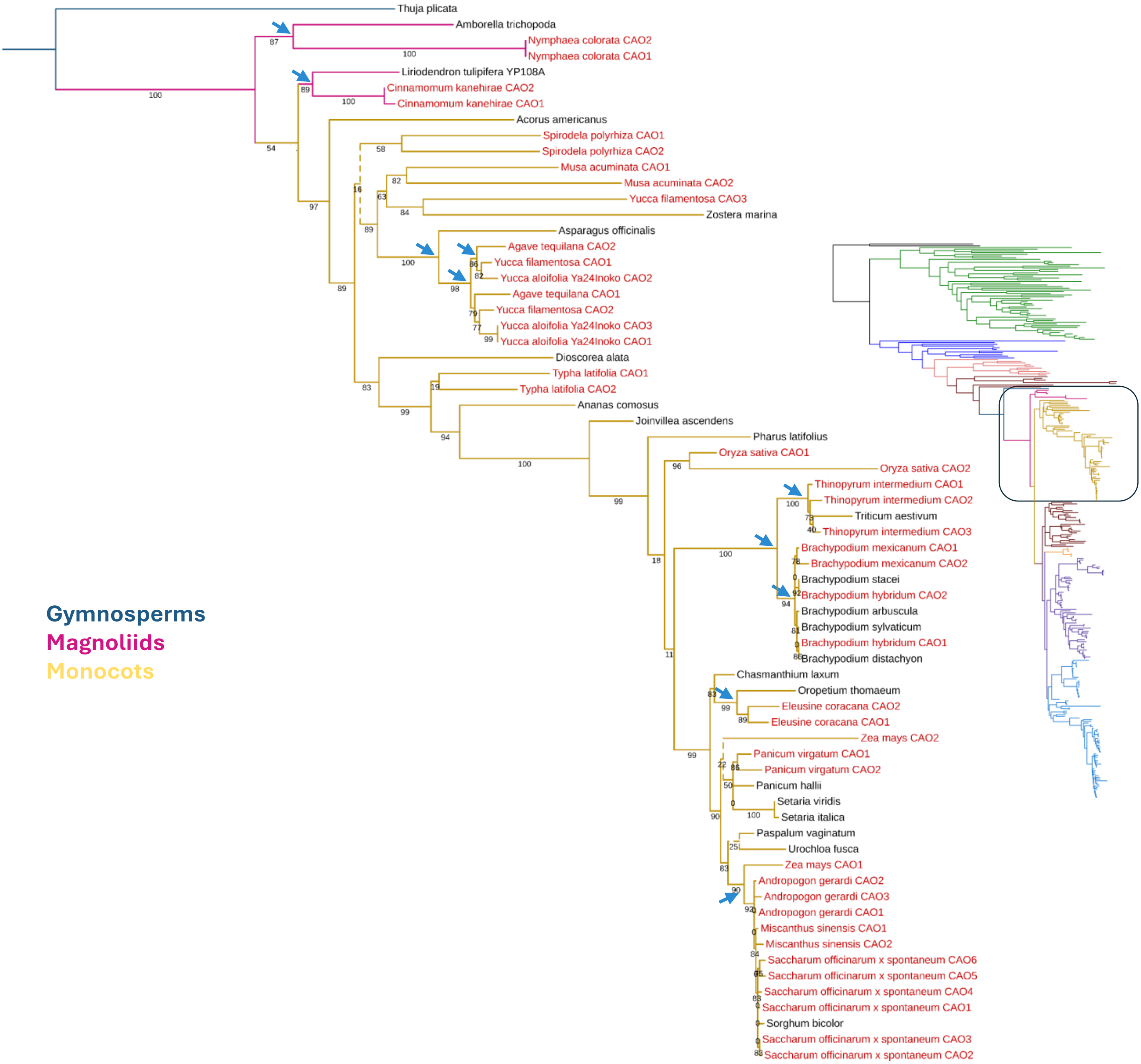


**Supplementary Figure 2**: (C) Gymnosperms, magnoliids, and monocots. (Continued)

D

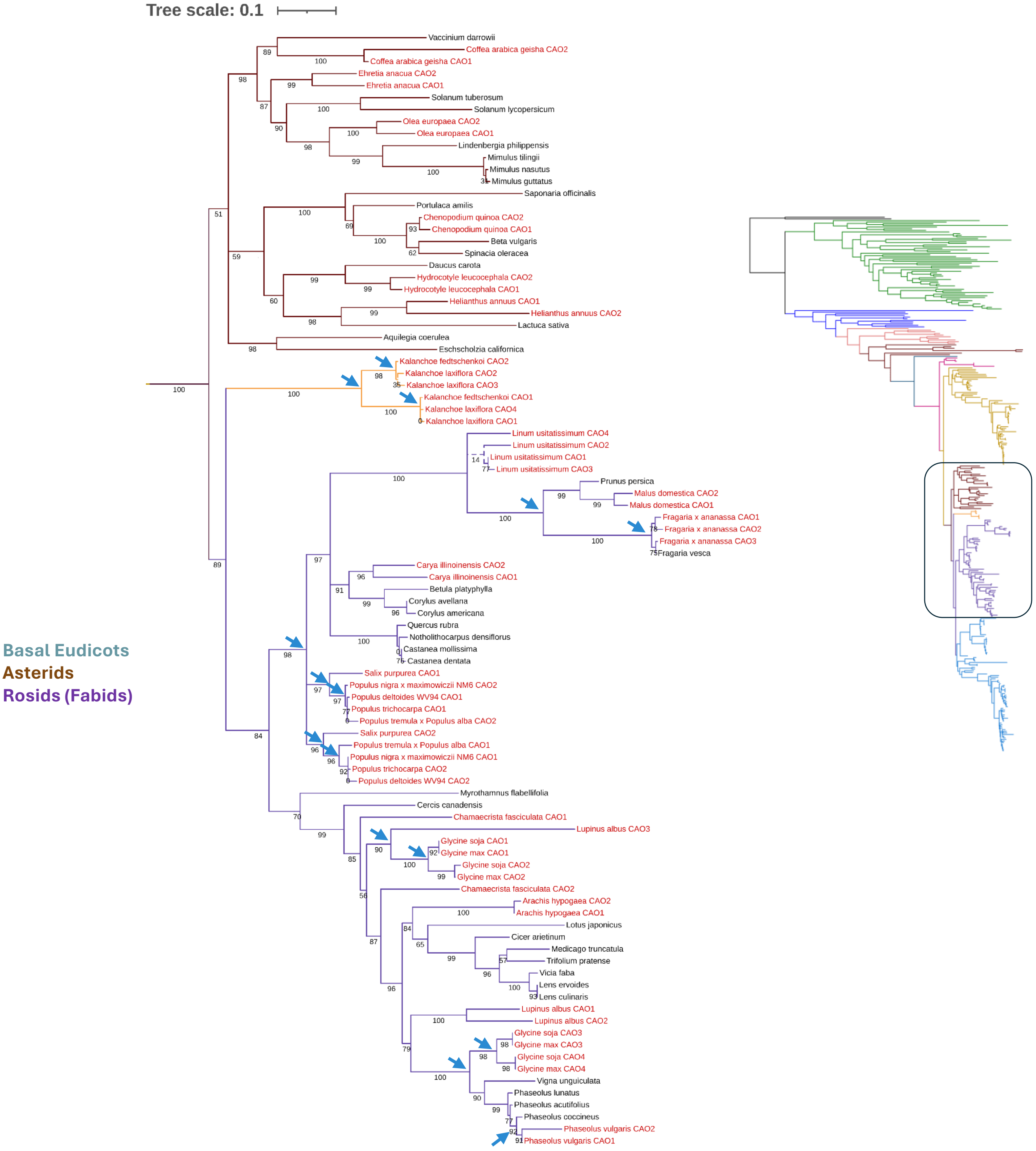


**Supplementary Figure 2**: (D) Basal eudicots, asterids and fabids. (Continued)

E

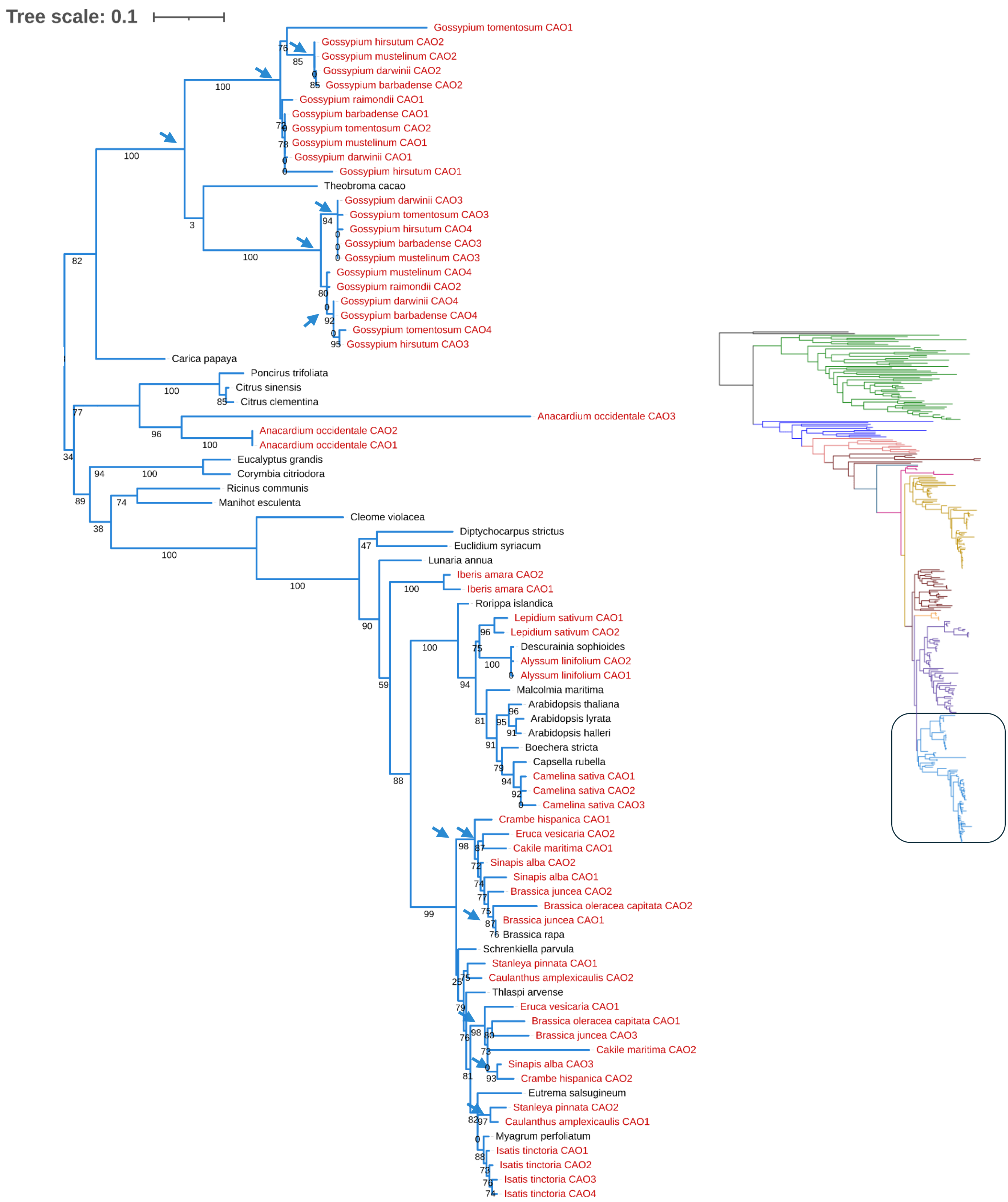


**Supplementary Figure 2:** (E) Malvids (Continued)

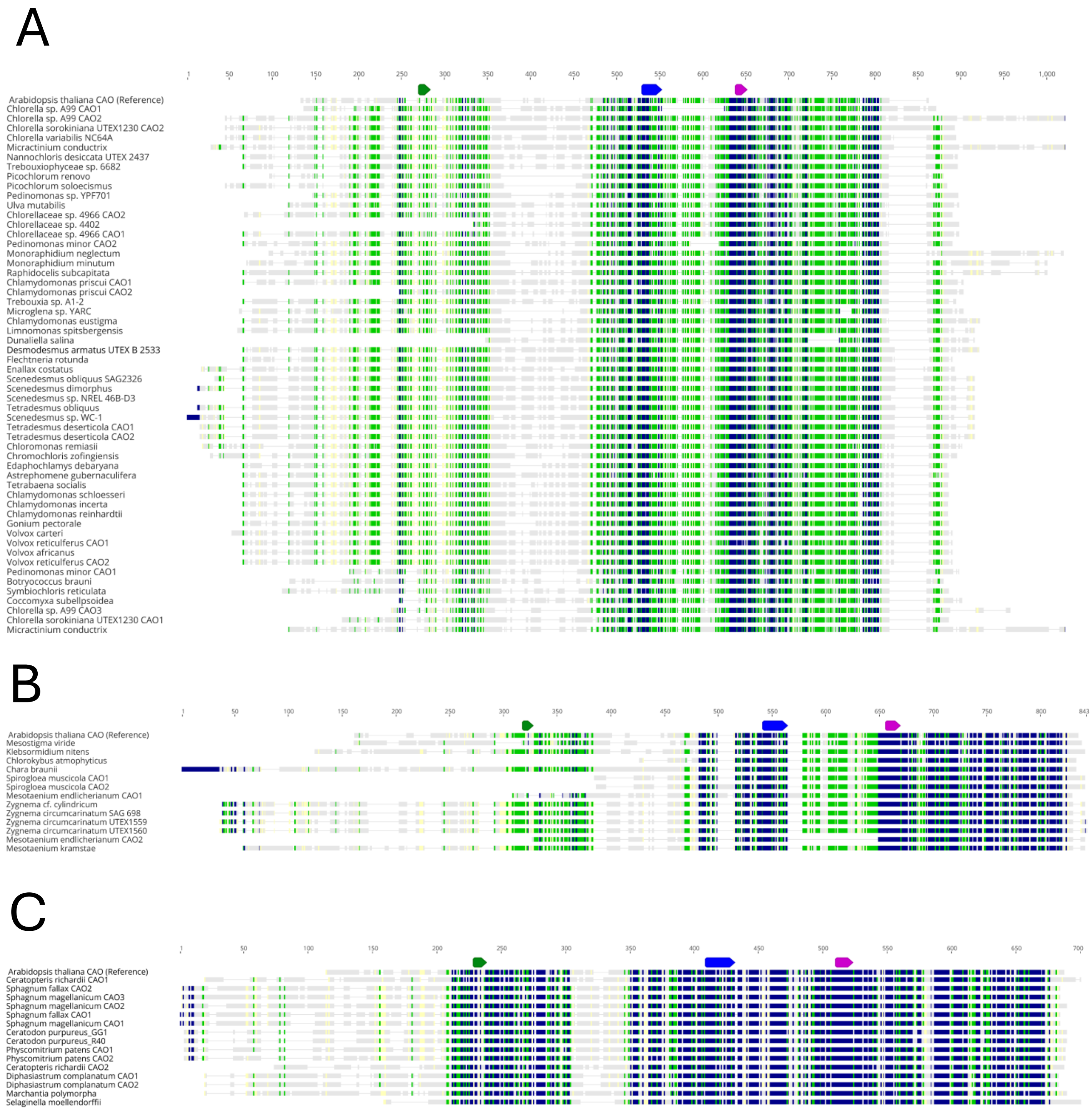


**Supplementary Figure 3**: Multiple sequence alignment of the predicted CAO amino acid sequences from Chlorophytes (A), Charophytes (B), non-vascular plants (C) and vascular plants (D). Global alignments were performed with MUSCLE. The colour of the residue depends on the degree of conservation among species (blue, fully conserved; green, 80–100% similar; yellow, 60–80% similar; gray, < 60% similar). As a reference, the CAO sequence from *Arabidopsis thaliana* is present in all alignments (top), with the key domains annotated (green, Degron sequence; blue, conserved Rieske cluster; purple, Fe-binding domain). Continued on the next page.

D

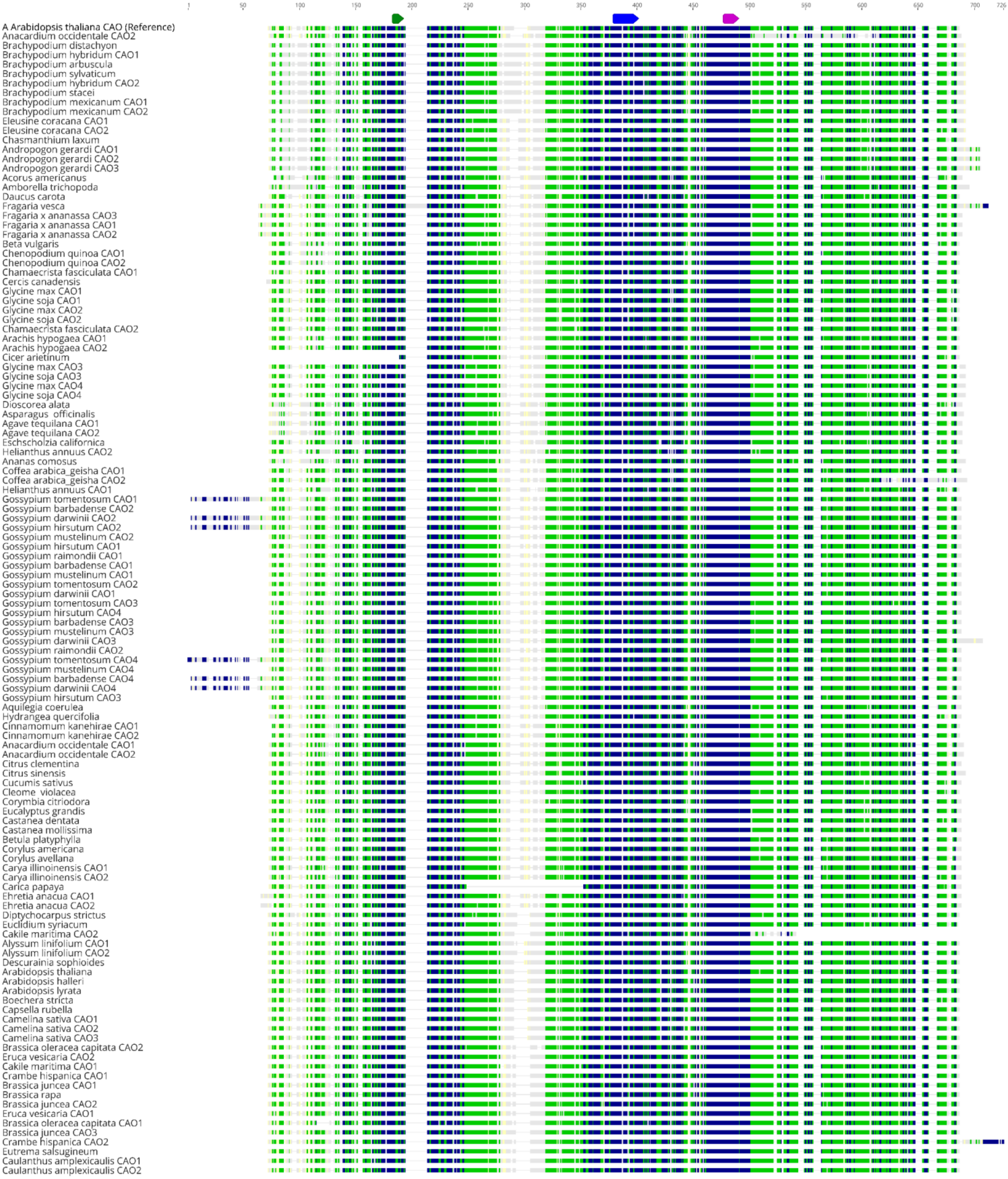


(Continued on the next page)

D

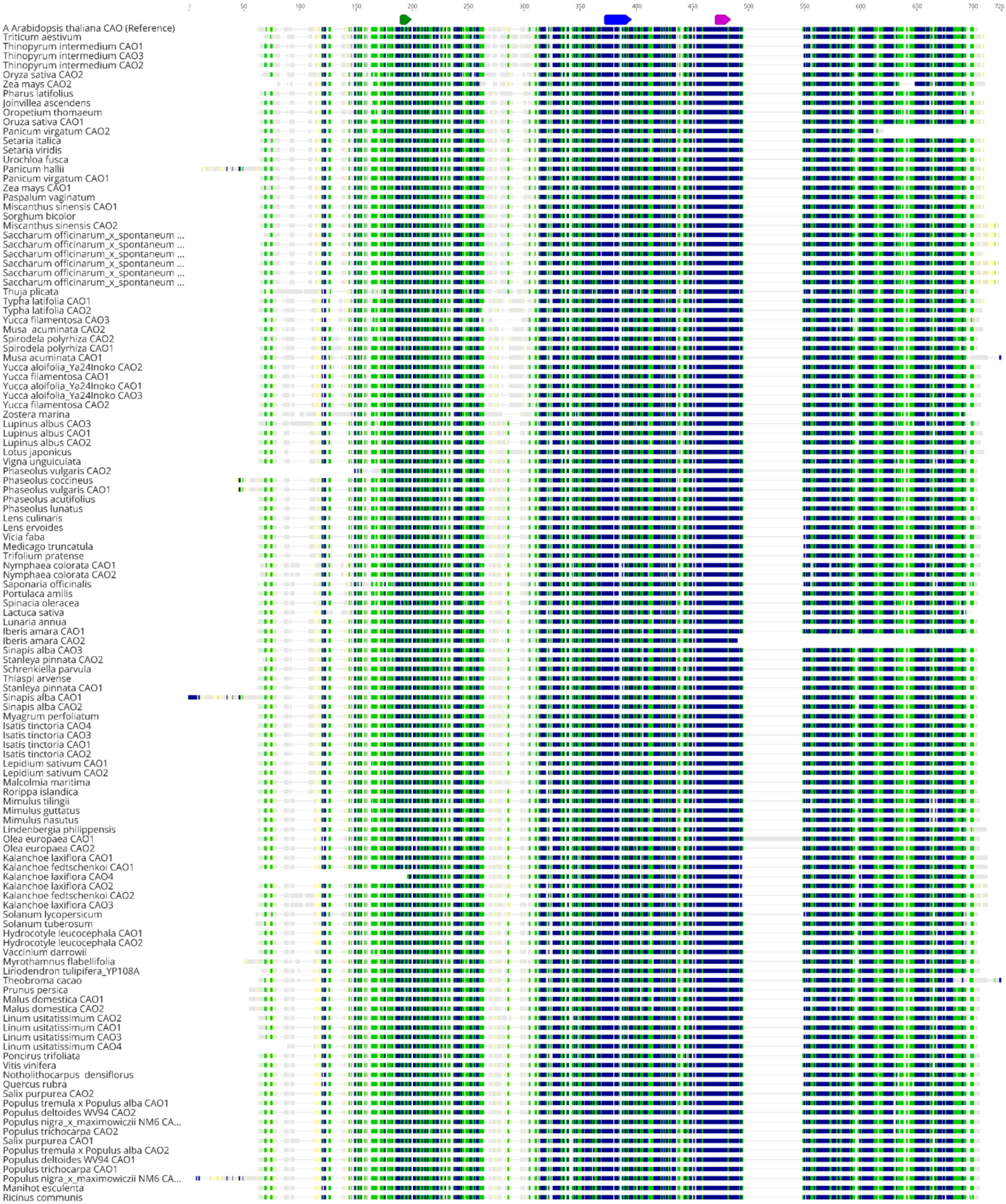
1
50
100
150
200
250
300
350
400
450
500
550
600
650
700
726
A Arabidopsis thaliana CAO (Reference)
Triticum aestivum
Thinopyrum intermedium CAO1
Thinopyrum intermedium CAO3
Thinopyrum intermedium CAO2
Oryza sativa CAO2
Zea mays CAO2
Pharus latifolius
Joinvillea ascendens
Oropetium thomaeum
Oruza sativa CAO1
Panicum virgatum CAO2
Setaria italica
Setaria viridis
Urochloa fusca
Panicum hallii
Panicum virgatum CAO1
Zea mays CAO1
Paspalum vaginatum
Miscanthus sinensis CAO1
Sorghum bicolor
Miscanthus sinensis CAO2
Saccharum officinarum_x_spontaneum ...
Saccharum officinarum_x_spontaneum ...
Saccharum officinarum_x_spontaneum ...
Saccharum officinarum_x_spontaneum ...
Saccharum officinarum_x_spontaneum ...
Saccharum officinarum_x_spontaneum ...
Thuja plicata
Typha latifolia CAO1
Typha latifolia CAO2
Yucca filamentosa CAO3
Musa acuminata CAO2
Spirodela polyrhiza CAO2
Spirodela polyrhiza CAO1
Musa acuminata CAO1
Yucca aloifolia_Ya24Inoko CAO2
Yucca filamentosa CAO1
Yucca aloifolia_Ya24Inoko CAO1
Yucca aloifolia_Ya24Inoko CAO3
Yucca filamentosa CAO2
Zostera marina
Lupinus albus CAO3
Lupinus albus CAO1
Lupinus albus CAO2
Lotus japonicus
Vigna unguiculata
Phaseolus vulgaris CAO2
Phaseolus coccineus
Phaseolus vulgaris CAO1
Phaseolus acutifolius
Phaseolus lunatus
Lens culinaris
Lens ervoides
Vicia faba
Medicago truncatula
Trifolium pratense
Nymphaea colorata CAO1
Nymphaea colorata CAO2
Saponaria officinalis
Portulaca amilis
Spinacia oleracea
Lactuca sativa
Lunaria annua
Iberis amara CAO1
Iberis amara CAO2
Sinapis alba CAO3
Stanleya pinnata CAO2
Schrenkiella parvula
Thlaspi arvense
Stanleya pinnata CAO1
Sinapis alba CAO1
Sinapis alba CAO2
Myagrum perfoliatum
Isatis tinctoria CAO4
Isatis tinctoria CAO3
Isatis tinctoria CAO1
Isatis tinctoria CAO2
Lepidium sativum CAO1
Lepidium sativum CAO2
Malcolmia maritima
Rorippa islandica
Mimulus tilingii
Mimulus guttatus
Mimulus nasutus
Lindenbergia philippensis
Olea europaea CAO1
Olea europaea CAO2
Kalanchoe laxiflora CAO1
Kalanchoe fedtschenkoi CAO1
Kalanchoe laxiflora CAO4
Kalanchoe laxiflora CAO2
Kalanchoe fedtschenkoi CAO2
Kalanchoe laxiflora CAO3
Solanum lycopersicum
Solanum tuberosum
Hydrocotyle leucocephala CAO1
Hydrocotyle leucocephala CAO2
Vaccinium darrowii
Myrothamnus flabellifolia
Liriodendron tulipifera_YP108A
Theobroma cacao
Prunus persica
Malus domestica CAO1
Malus domestica CAO2
Linum usitatissimum CAO2
Linum usitatissimum CAO1
Linum usitatissimum CAO3
Linum usitatissimum CAO4
Poncirus trifoliata
Vitis vinifera
Notholithocarpus densiflorus
Quercus rubra
Salix purpurea CAO2
Populus tremula x Populus alba CAO1
Populus deltoides WV94 CAO2
Populus nigra_x_maximowiczii NM6 CA...
Populus trichocarpa CAO2
Salix purpurea CAO1
Populus tremula x Populus alba CAO2
Populus deltoides WV94 CAO1
Populus trichocarpa CAO1
Populus nigra_x_maximowiczii NM6 CA...
Manihot esculenta
Ricinus communis
